# Genome-wide quantification of the effects of DNA methylation on human gene regulation

**DOI:** 10.1101/146829

**Authors:** Amanda J. Lea, Christopher M. Vockley, Rachel A. Johnston, Christina A. Del Carpio, Luis B. Barreiro, Timothy E. Reddy, Jenny Tung

## Abstract

Changes in DNA methylation are important in development and disease, but not all regulatory elements act in a methylation-dependent (MD) manner. Here, we developed mSTARR-seq, a high-throughput approach to quantify the effects of DNA methylation on regulatory element function. We assay MD activity in 14% of the euchromatic human genome, identify 2,143 MD regulatory elements, and predict MD activity using sequence and chromatin state information. We identify transcription factors associated with higher activity in unmethylated or methylated states, including an association between pioneer transcription factors and methylated DNA. Finally, we use mSTARR-seq to predict DNA methylation-gene expression correlations in primary cells. Our findings provide a map of MD regulatory activity across the human genome, facilitating interpretation of the many emerging associations between methylation and trait variation.

## Main text

DNA methylation—the covalent addition of methyl groups to nucleotide bases, most often at CpG motifs—is a gene regulatory mechanism that plays a fundamental role in development, disease susceptibility, and the response to environmental conditions^1–6^. These functions suggest that variation in DNA methylation should be important in explaining trait variation. In support of this idea, epigenome-wide association studies (EWAS) have now identified thousands of statistical relationships between phenotypic variation and DNA methylation levels at individual CpG sites across the genome^7^.

However, not all changes in DNA methylation causally affect gene regulation^8,9^, making variation in DNA methylation more functionally important at some loci than others. Mapping methylation-dependent (MD) regulatory activity across the genome is therefore essential for interpreting the growing number of DNA methylation-trait associations, as well as understanding the basic biology of epigenetic gene regulation. Current approaches for assaying MD activity are either too low-throughput to support genome-scale analyses or have focused on measuring methylation-dependent transcription factor binding outside the cellular context^8,10–17^ (Table S1). These studies suggest widespread differential TF sensitivity to DNA methylation levels^15–17^, but leave open whether, and to what degree, differential sensitivity translates to differences in gene expression itself.

To address these questions, we developed a high-throughput method, mSTARR-seq, that assays the causal relationship between DNA methylation and regulatory activity within a cellular context. mSTARR-seq combines genome-scale strategies for quantifying enhancer activity via self-transcribing episomal reporter assays (e.g., STARR-seq^18^) with enzymatic manipulation of DNA methylation at millions of unique CpG sites (Fig. 1). To eliminate the confounding effects of DNA methylation in the vector itself, we engineered a CpG-free mSTARR-seq-specific vector (*pmSTARRseq*) that also eliminates the potential for bacterial *Dam-* or *Dcm*-mediated methylation (Fig. 1A). As in STARR-seq, the *pmSTARRseq* vector enables a library of query fragments to be inserted in the 3’ untranslated region of a constitutively expressed reporter gene, such that fragments with regulatory activity drive their own transcription when transfected into a cell type of interest^18^. Prior to transfection, the plasmid input library can be treated with either the methyltransferase *M.SssI*, which methylates all CpG sites, or a sham treatment, which leaves them unmethylated. The regulatory activity of fully methylated fragments can then be compared to the activity of unmethylated fragments by using high-throughput sequencing to quantify their relative abundances in reporter gene-derived mRNA (Fig. 1B).

**Figure 1.**
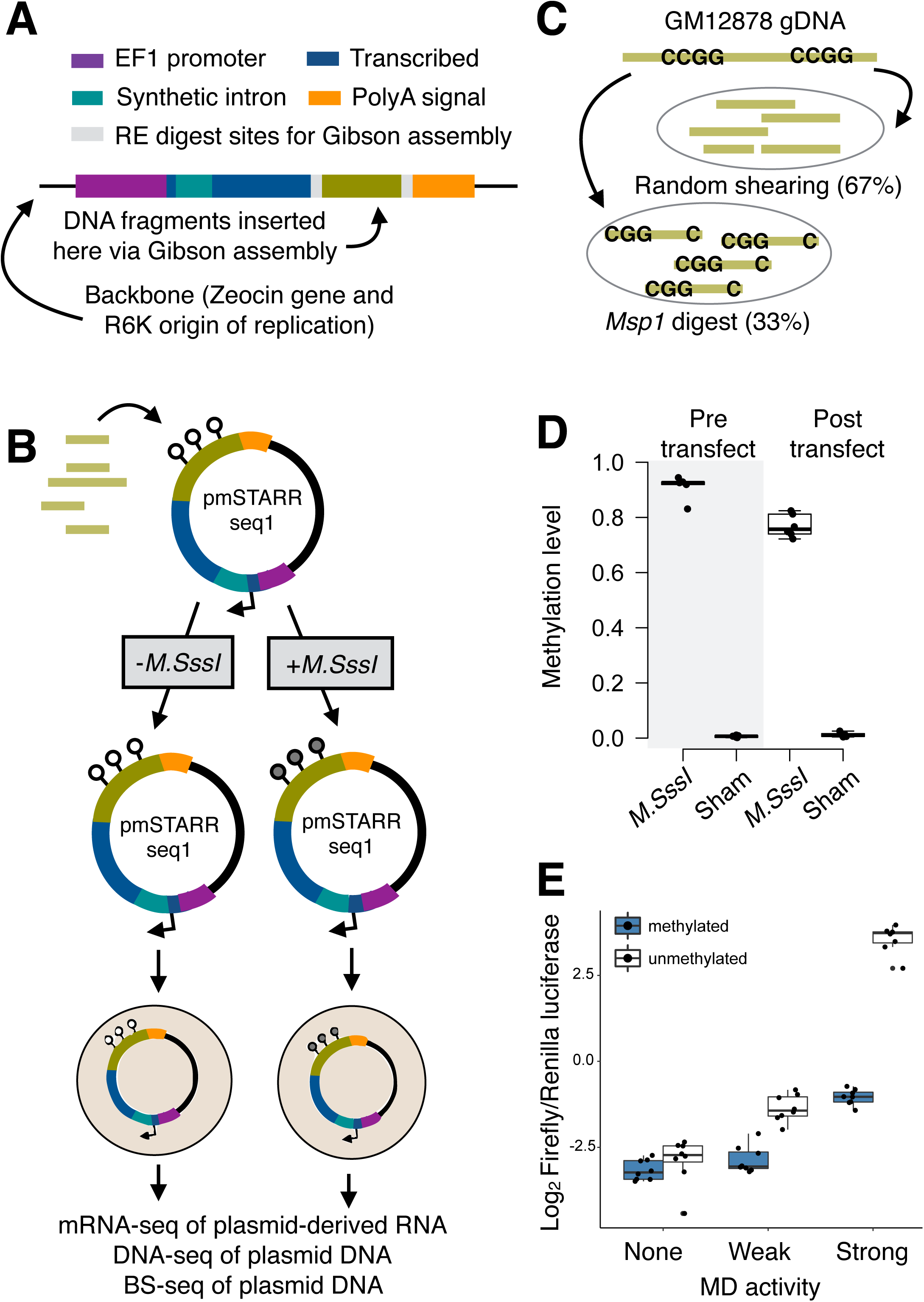
mSTARR-seq experimental design. (A) The *pmSTARRseq* vector is entirely CpG free. It is designed so that functional regulatory elements will self-transcribe to produce a fully processed mRNA transcript, including a transcribed region (dark blue) that spans a synthetic intron (teal), the sequence of the regulatory element itself (green), and an SV40 polyA signal (orange). (B) DNA fragments are cloned into *pmSTARRseq* in high-throughput. The resulting library is subjected to either experimental methylation (*M.SssI* treatment) or a sham treatment, and each pool is transfected into a cell line of interest (here, we used the K562 myeloid cell line; n=6 replicates per condition). After a 48 hr incubation period, plasmid DNA and plasmid-derived mRNA are extracted and the variable insert regions sequenced. (C) As input, we used GM12878 DNA fragmented through random shearing or *Msp1* digest (to enrich for CpG-containing regions of the genome). The resulting fragment pools were mixed in a 2:1 ratio. (D) Bisulfite sequencing of the GM12878 plasmid pool pre- and post-transfection confirms that *M.SssI* treatment almost completely methylates CpG sites contained in the candidate regulatory elements. High methylation levels are maintained throughout the experiment. Y-axis shows mean CpG methylation level per experimental replicate. (E) Low-throughput validation (CpG-free luciferase reporter assay^12^) of three candidate regulatory elements with no (FDR>0.2), weak (0.05<FDR<0.1), or strong evidence (FDR<0.001) for MD activity in mSTARR-seq (Wilcoxon p-value, comparison between conditions: 0.069, 1.55x10^−4^, and 1.55x10^−4^, respectively).

To quantify MD activity across the human genome, we combined *MspI*-digested genomic DNA (to enrich for CpG-containing fragments) with randomly sheared DNA from the HapMap GM12878 cell line (Fig. 1C). We then transfected unmethylated and methylated versions of the plasmid library (n=6 replicates per condition) into the K562 cell line. Forty-eight hours post-transfection, we isolated and sequenced both the plasmid-derived mRNA and the fragment inserts from each plasmid DNA pool (Table S2; fig. S1). We also performed bisulfite sequencing on the plasmid DNA to confirm maintenance of the expected DNA methylation state throughout the experiment (Fig. 1D).

In total, we assayed ~750,000 unique DNA fragments in each library (mean ± SD = 759,725 ± 252,187 fragments per replicate; one replicate from the methylated condition was excluded from all analyses due to low sequencing depth), comparable to or exceeding the diversity in published STARR-seq and massively parallel reporter assays (fig. S2). For subsequent analysis, we binned the genome into 200 bp non-overlapping intervals and filtered these regions to focus on the 277,896 intervals that overlapped at least 1 mRNA read and 1 DNA read in at least half of the replicates in each condition. These 277,896 intervals were covered by 724,391 unique fragments of size 314 bp ± 105 bp (mean ± S.D.; fig. S3). This stringently filtered data set represents 1.83 million unique CpG sites, 57% of fragments expected from a complete *MspI* digest of the human genome, and 14% of the euchromatic genome of the K562 cell line (fig. S4).

We first focused on regions with regulatory capacity (i.e., enhancer-like activity), whether in the unmethylated condition, methylated condition, or both. We identified 24,945 intervals of 200 bp (9% of analyzed regions, at a 10% false discovery rate) in which the abundance of plasmid-derived mRNA was significantly greater than the amount of input plasmid DNA (Table S3). As expected, the set of regions capable of enhancer-like activity was highly enriched for K562 ENCODE chromatin states^19^ associated with H3K4me1 and H3K27ac, which mark active enhancers (Fisher’s exact test, log_2_ odds=2.53, p<10^−15^) and highly depleted in regions that lacked both marks (log_2_ odds=-0.94, p<10^−15^; Fig. 2A). Regions that overlapped H3K4me1 and H3K27ac-marked chromatin states also consistently displayed the largest effect sizes (relative to regions that lacked these marks, or only exhibited one mark; linear model, p<10^−15^; Fig. 2B). Finally, regions annotated as strong enhancers in K562 cells exhibited the strongest effects of all 12 chromatin states (p<10^−15^), and contained the largest proportion of elements with significant regulatory activity relative to any other chromatin state (at a 10% FDR, 37% of regions tested had significant activity). In general, power to detect enhancer activity increased with larger query fragment sizes (Fig. 2C), suggesting that short fragments may eliminate binding sites key to functional enhancer activity.

**Figure 2.**
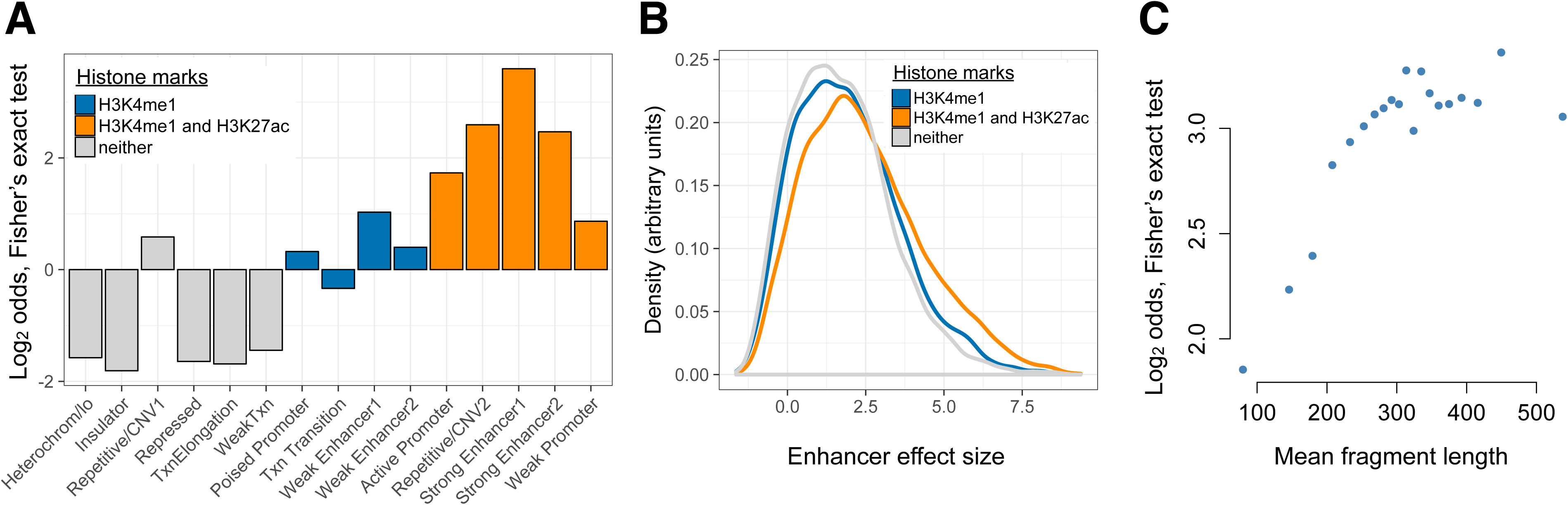
mSTARR-seq identifies regions with endogenous regulatory activity. (A) Regions with significant regulatory activity in the mSTARR-seq assay are enriched for chromatin state annotations defined by active marks (H3K4me1 and H3K27ac, colored orange). The y-axis depicts the log_2_(odds) from a two-sided Fisher’s exact test for enrichment (or depletion) of mSTARR-seq identified enhancers in each of the 12 annotated chromatin states in K562 cells (p<0.05 for all tests). Positive y-axis values indicate enrichment and negative values indicate depletion. (B) Effect sizes for loci with significant enhancer activity (FDR<10%; x-axis) are consistently larger for mSTARR-seq identified enhancers that occur in chromatin state annotations defined by active marks. (C) Binning regions with significant mSTARR-seq enhancer activity by fragment length reveals that larger fragments are more strongly enriched for ENCODE-annotated ‘strong enhancers’. The y-axis depicts the log_2_(odds) from a Fisher’s exact test for enrichment of mSTARR-seq enhancers (binned by deciles of fragment length) in either of the two ‘strong enhancer’ chromatin states (p<0.05 for all tests).

We next investigated which regulatory elements were functionally affected by DNA methylation marks. We identified 2,143 regions with significant MD activity (8.59% of those tested; 10% FDR), 88% of which were more active when unmethylated and 12% which were more active when methylated (Fig. 3A; Table S4). Only 4 of the 941 CpG-free regions in the analysis set (0.4%) were inferred to have MD activity, indicating a low false positive rate (Fig. 3B). Estimates of MD activity from mSTARR-seq were also consistent with estimates from single-locus luciferase reporter assays^13^ (Fig. 1E). Overall, we found that MD enhancers have higher CpG densities and contain more CpG sites than non-MD enhancers (Wilcoxon-signed rank test, W=3.51x10^7^, p<10^−15^; Fig. 3C). However, CpG density only explained 6.8% of variation in the magnitude of methylation dependence, suggesting that other characteristics also contribute to quantitative variation in MD activity (Spearman’s rho=0.246, p<10^−15^; Fig. 3D).

**Figure 3.**
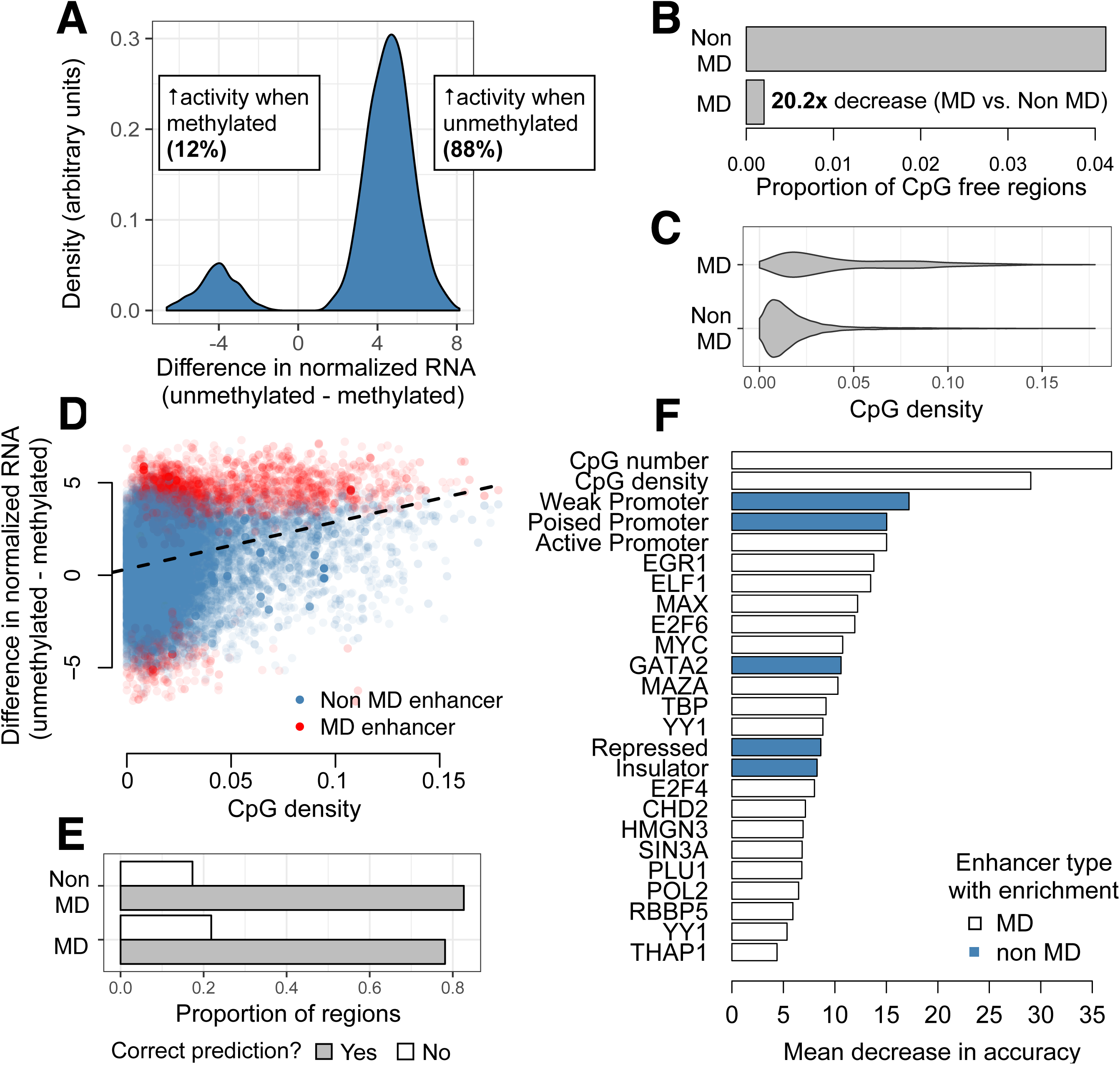
mSTARR-seq identification and prediction of MD enhancers. (A) The distribution of differences in normalized mRNA transcript abundance between the unmethylated and methylated conditions (all significant MD enhancers are shown). (B) CpG-free MD enhancers occur at a 20.2-fold lower rate than CpG-free windows with no MD enhancer activity. (C) Distribution of fragment CpG density for regions identified as MD versus non-MD enhancers. (D) CpG-dense mSTARR-seq enhancers tend to be repressed by DNA methylation, such that mRNA abundance is higher in the unmethylated condition relative to the methylated condition (positive y-axis value). X-axis: CpG sites/fragment window length (Spearman’s rho for correlation between x and y axes=0.246, p<10^−15^; n=24,945 regions with significant regulatory element activity). (E) The proportion of non-MD and MD enhancers that were accurately classified via a random forests (RF) classifier. (F) Features that distinguish MD and non-MD enhancers in the RF classifier (10% FDR). X-axis: mean decrease in predictive accuracy when excluding the focal variable. Blue: positive prediction of non-MD enhancers; white: positive prediction of MD enhancers.

To explore these characteristics, we used a random forests classifier to evaluate the contribution of 147 genomic features to differentiating MD enhancers (specifically, the n=1866 regions suppressed by methylation) from non-MD enhancers (n=5703 regions that exceed an FDR of 50% in our test for MD activity). Our feature set included information about CpG site density; endogenous chromatin state, chromatin accessibility, and DNA methylation levels^19,20^; evolutionary conservation^21^; and TF binding from K562 ENCODE ChIP-seq data^19^ (Table S5). The resulting RF model predicted MD regulatory element activity with 82% accuracy (Fig. 3E). In addition to CpG site information, 25 features were identified as key predictors based on two measures of variable importance, the mean decrease in accuracy and the Gini coefficient (FDR<10%; Fig. 3 and Table S5). Relative to non-MD enhancers, enhancers suppressed by DNA methylation were more likely to occur in regions with endogenous promoter activity and less likely to occur in endogenously repressed regions of the genome. MD enhancers were also more likely to contain binding sites for the TFs ELF1, E2F6, MAX, and MYC, all of which have CpG sites in their canonical binding motifs (Fig. 3F).

Previous work indicates that many TFs are sensitive to DNA methylation levels in or near their binding motifs^15–17^. This ability to “read” epigenetic modifications to DNA sequence could explain, at least in part, variation in MD regulatory activity in our data set. Indeed, among the 1866 MD enhancers in which DNA methylation suppresses activity, we identified 24 significantly enriched TF binding motifs (relative to the background set of all regions with mSTARR-seq regulatory activity; 1% FDR). 15 of these motifs belong to the ETS family, a 6.6x enrichment over chance (hypergeometric test p=3x10^−13^; Fig. 4A and Table S6). ETS binding is thought to be methylation dependent for ‘Class I’ ETS TFs^22–27^, which bind the canonical motif ACCGGAAGT, but not for ‘Class III’ ETS family TFs, whose binding motifs do not consistently include CpG sites^28^. In support, 12 of the 15 ETS TFs we identified belong to Class I, and none belong to Class III. The remaining 3 belong to Class II, for which methylation-dependent binding was previously unexplored: our results suggest they behave more similarly to Class I than Class III.

**Figure 4.**
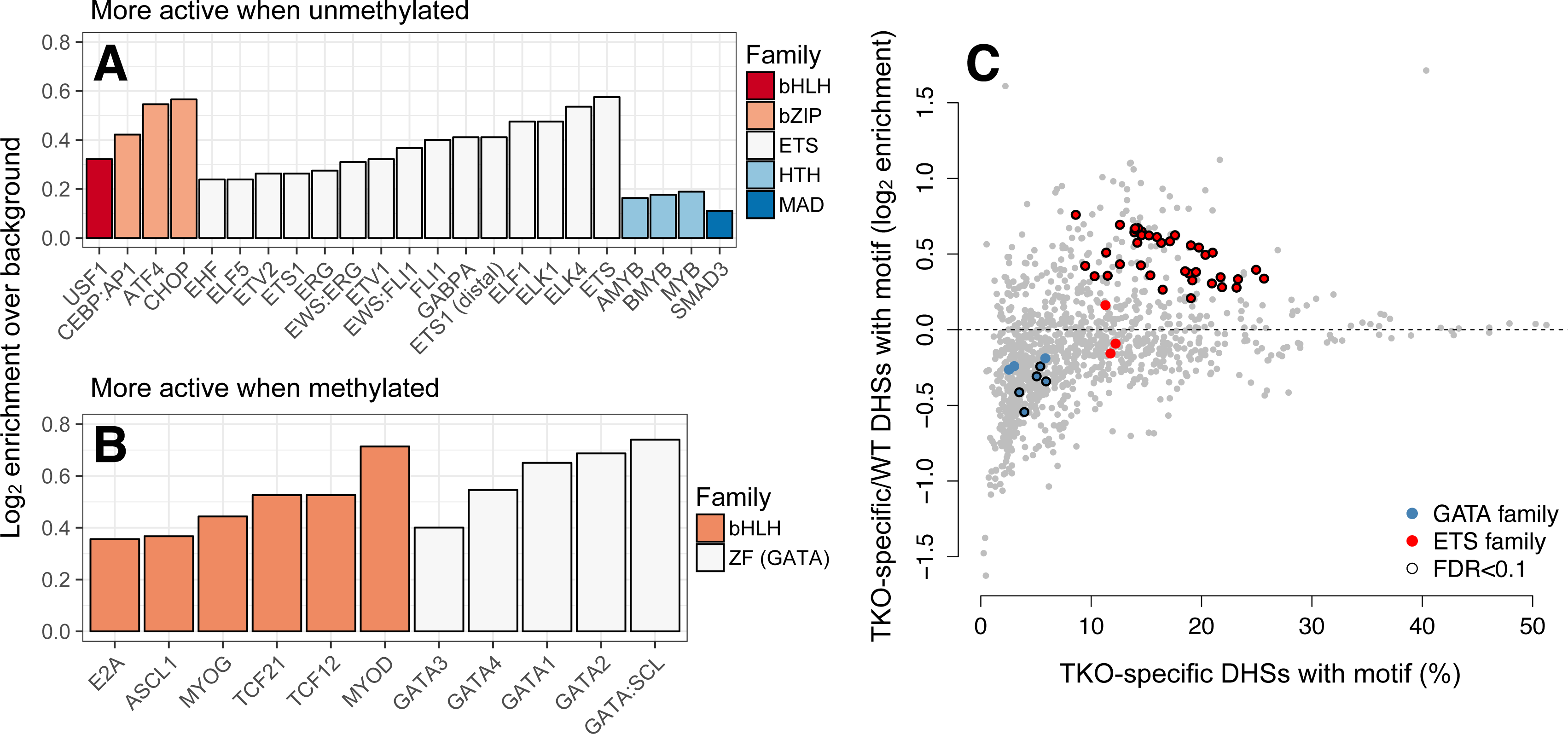
mSTARR-seq identifies MD-dependent transcription factor-DNA binding. (A) Transcription factor motifs that are enriched in MD enhancers that are more active when unmethylated, colored by TF family. (B) TF motifs that are enriched in MD enhancers that are more active when methylated. (C) DNase hypersensitive sites (DHS) specific to murine stem cells that lack DNA methylation (DNMT triple knock-outs: TKO) are strongly enriched for ETS family binding sites relative to wild type cells with intact DNA methylation. In contrast, DHSs specific to wild type cells are enriched for GATA family binding sites relative to triple knockouts. DHS data are from^25^. X-axis: percent of knockout-specific DHSs that contain a given TF binding motif (n=1251 motifs). Y-axis: Ratio of knockout versus wild-type specific DHSs containing a given TF binding site motif. Colored dots circled in black show significant enrichment for a ETS or GATA family TF (10% FDR in a hypogeometric test).

We also identified 9 significantly enriched TF binding motifs in the 257 MD enhancers with increased activity in the methylated condition (1% FDR). TFs from the basic helix-loop-helix (bHLH) family and GATA subfamily of zinc finger TFs were strongly enriched in this set (a 2.91x and 20x enrichment over chance, hypergeometric test p=0.33 and p=1.99x10^−7^, respectively; Fig. 4B and Table S7), consistent with reports that GATA3, GATA4, and bHLH family TFs bind to methylated DNA outside the cellular context^16^. We compared our findings to published chromatin accessibility data for wild type murine stem cells, which contain normal patterns of DNA methylation, and triple knockouts for *DNMT1, DNMT3a*, and *DNMT3b*, in which DNA methylation is abolished^29^. For 5 of 10 tested GATA family TFs, open chromatin regions specific to wild type (i.e., those absent in the triple knockouts) were significantly enriched for their cognate binding sites (Fig. 4C), in support of the idea that GATA family TFs preferentially bind methylated DNA in service of their function as “pioneer” factors^30^. In contrast, ETS family TF binding sites were almost universally (38 of 41 tested) enriched in DNMT knockout-specific open chromatin regions.

Finally, for mSTARR-seq results to be maximally useful in interpreting DNA methylation-trait associations, we reasoned that they should explain the substantial heterogeneity in DNA methylation-gene expression correlations observed in real populations. To test this possibility, we drew on paired DNA methylation and gene expression data for 1202 human primary monocytes^31^ (a cell type closely related to K562s), in which the mean correlation between DNA methylation levels and gene expression at the nearest gene is 0.006 +/- 0.189 s.d. (and -0.023 +/- 0.304 for CpG sites significantly (FDR<10%) correlated with gene expression; n=81,883 site-gene pairs). Genome-wide, we observed that significant DNA methylation-gene expression correlations in monocytes (FDR<10%) were moderately enriched in mSTARR-seq MD enhancers versus non-MD enhancers (Fisher’s exact test, log_2_ odds=0.60, p=3.38x10^−4^). However, for CpG sites that display the canonical negative correlation between DNA methylation and gene expression levels, this relationship was greatly strengthened (log_2_ odds=1.02, p<10^−15^). Thus, mSTARR-seq can identify the CpG sites for which DNA methylation variation is most tightly linked to gene expression variation in human primary cells.

Together, our findings emphasize substantial variability in the functional relationship between DNA methylation and gene regulation across the genome. Using mSTARR-seq, we show that the magnitude of this relationship is both predictable from genome characteristics and in turn predicts *in vivo* heterogeneity in real populations. The resulting map of MD regulatory activity thus provides useful guidance for prioritizing DNA methylation-trait associations for further investigation: CpG sites in which DNA methylation levels causally influence gene expression are more likely to be of interest than those that are effectively silent. In addition, we provide support for the hypothesis that pioneer TFs, such as members of the GATA TF family, have a higher affinity for methylated DNA, potentially aiding in their ability to bind condensed chromatin^30^. Indeed, in addition to GATA family TFs, TFs important in development and cell fate, such as FOXA, MyoD, and TCF21, are enriched among MD enhancers with increased activity when methylated. These results raise the interesting possibility that preferential binding of methylated loci could be used to aid in pioneer TF discovery. Finally, mSTARR-seq can be applied as an efficient, high-throughput strategy to map MD activity in a variety of settings, including at specific loci of interest, across cell types, or across cellular environments. Epigenome editing approaches will be useful for following up the most interesting loci.

## ACKNOWLEDGMENTS

We thank Michael Yuan and members of the Reddy and Tung labs for experimental contributions and helpful discussions, and the Rehli lab for the gift of the pCpGL vector. This work was supported by a Sloan Foundation Early Career Research Fellowship, NIH grants R01-GM102562 and R21-AG049936, and NSF grant BCS-1455808. RNA-seq and DNA-seq data will be deposited in NCBI’s Short Read Archive following publication. Code and data will be available at https://github.com/AmandaJLea/mstarr_seq following publication. The mSTARR-seq protocol is available online at www.tung-lab.org/protocols-and-software.html. The *pmSTARRseq* vector and DNA input library used in the experiments described here (fig. S5) can be shared with interested third parties pending a mutual transfer and non-commercial use agreement (see details at www.tung-lab.org/protocols-and-software.html).

## Supplementary Materials

### Author Contributions

**Materials and Methods**
*Laboratory techniques and methods*

Text S1. pmSTARRseq design
Text S2. Generation of plasmid libraries for mSTARR-seq
Text S3. Cell culture, plasmid transfection, and cell harvesting
Text S4. Isolation and preparation of mRNA derived from the mSTARR-seq plasmid
Text S5. Preparation of plasmid DNA for DNA-seq and bisulfite sequencing
Text S6. Luciferase reporter assays
*Computational techniques and methods*

Text S7. Low-level data processing
Text S8. Identification of enhancers and methylation dependent (MD) enhancers
Text S9. Annotation of analyzed mSTARR-seq fragments
Text S10. *In silico MspI* digest
Text S11. Random forests classification
Text S12. Transcription factor binding motif enrichment analyses
Text S13. Correlations between DNA methylation and gene expression levels in primary cells
**Figures S1-S5**

Figure S1. Diversity in plasmid DNA-seq libraries versus mRNA-seq libraries.
Figure S2. Fragment diversity in mSTARR-seq experiments versus other published
multiplexed reporter assays (MPRA) or STARR-seq experiments.
Figure S3. Distribution of analyzed fragment lengths.
Figure S4. Regions covered by mSTARR-seq.
Figure S5. Retransforming a plasmid pool results in almost no loss in diversity.
**Tables S1-S8**

Table S1: Other methods for testing the causal relationship between DNA methylation and gene regulation.
Table S2: Samples sequenced in this study.
Table S3: Linear model results testing for mSTARR-seq regulatory activity.
Table S4: Linear model results testing for methylation-dependent regulatory activity.
Table S5: Random forests analysis results.
Table S6: TF motif enrichment results for MD enhancers with greater activity in the unmethylated condition.
Table S7: TF motif enrichment results for MD enhancers with greater activity in the methylated condition.
Table S8A: Luciferase reporter assay details.
Table S8B: Luciferase reporter assay results.

